# Depth-resolved fiber photometry of amyloid plaque signals in freely behaving Alzheimer’s disease mice

**DOI:** 10.1101/2025.04.08.647900

**Authors:** Nicole Byron, Niall McAlinden, Filippo Pisano, Marco Pisanello, Jacques Ferreira, Cinzia Montinaro, Keith Mathieson, Massimo De Vittorio, Ferruccio Pisanello, Shuzo Sakata

## Abstract

Alzheimer’s disease pathology typically manifests itself across multiple brain regions yet assessment at this scale in mouse models remains a challenge. This hinders the development of novel therapeutic approaches. Here we introduce a novel fiber photometry approach to monitor amyloid pathology in freely behaving mice. We first demonstrated that flat fiber-based photometry can detect amyloid signals across multiple brain regions under anesthesia after injecting a blood-brain barrier permeable tracer, Methoxy-X04. The depth profile of in vivo fluorescent signals was correlated with postmortem histological plaque signals. After confirming its feasibility ex vivo, we chronically implanted a tapered fiber for depth-resolved fiber photometry in freely behaving mice. After injecting Methoxy-X04, fluorescent signals increased in a depth-specific manner in Alzheimer’s mice, but not in wild-type littermates. While fiber photometry has been widely adopted to monitor neuronal and non-neuronal activity, our approach expands the capabilities to monitor molecular pathologies such as amyloid plaques, even in a freely behaving condition.

## Introduction

Alzheimer’s disease (AD) is the most common form of dementia. Amyloid plaques have long been recognized as a hallmark of AD (1–3). Recent therapeutics targeting amyloid-β protofibrils or deposited amyloid plaques have proven effective in patients (4, 5), successful in translating early preclinical mouse data (6) to the clinic. As such, evaluating novel interventions in preclinical mouse models plays a critical role in accelerating further successes.

While a wide range of pharmacological and non-pharmacological intervention approaches have been examined in mouse models, postmortem biochemical and histological analysis remains the gold standard in the field. This means that every intervention is a one-time experiment that requires animal sacrifice. To optimize an intervention approach flexibly, it would be ideal to evaluate its efficacy in real-time.

Although non-invasive approaches to assess a pathological state are available for human subjects and plasma-based biomarker detection is an emerging area (7–9), options for mouse models remain limited. For example, although microdialysis allows assessment of interstitial fluid contents in the brain (10–12), it lacks spatial resolution. Two-photon imaging can detect plaques *in vivo* by combining with a blood-brain barrier permeable tracer, Methoxy-X04 (13–15). However, depth penetration is limited. It is crucial to assess the plaque distribution at depth, given the fact that amyloid pathology can be seen across multiple deep brain regions. Another approach is optoacoustic tomography (16), allowing brain-wide monitoring of plaque signals. However, current technologies do not permit monitoring of amyloid pathology across multiple brain regions in a freely behaving condition.

Fiber photometry is an alternative, versatile optical approach (17–20). It allows monitoring of neuronal and non-neuronal population activity in a cell-type-specific manner *in vivo* (21–25). While fiber photometry has also been adopted to monitor intracellular signaling (26), it remains unknown if fiber photometry can monitor extracellular pathological signals like amyloid plaques. For instance, fiber photometry typically uses fluorescent sensors that require availability and expression of its genetically encoded construct. To overcome this requirement, molecular pathologies could be assessed through a non-genetic strategy by administration of a blood-brain barrier fluorescent plaque dye.

Here we test the hypothesis that fiber photometry can be adapted to access molecular pathologies, using a non-genetic approach. Specifically, we test if it can be used to monitor amyloid pathology in 5xFAD mice, a widely used AD mouse model carrying five familial AD mutations (27). To this end, we take two approaches. First, as a proof-of-concept, we utilize conventional flat optical fibers to examine if amyloid pathology can be monitored across multiple depths in mice. Second, we exploit the photonic properties of tapered optical fibers (20) to establish depth-resolved photometry of plaque signals *ex vivo* and *in vivo*. Our novel photometry approach expands the capabilities of *in vivo* fiber photometry to examine the pathological features of an AD mouse model in a freely behaving condition.

## Materials and Methods

### Animals

Experiments were performed in accordance with the UK Animals (Scientific Procedures) Act of 1986 Home Office regulations and approved by the Home Office (PP0688944). Mice were housed with sex-matched littermates, if available, on a 12-hour/12-hour light/dark cycle with access to food and water ad libitum. All experiments were performed during the light period. 5xFAD mice (27) (JAX006554, The Jackson Laboratory) were obtained and backcrossed onto C57BL/6 background (>F10). Male and female 5xFAD+/-, referred to as 5xFAD, and 5xFAD-/-, referred to as WT, mice aged 3-12 months were used (**Supplementary Table 1**). 15 mice (7 5xFAD, 6 males and 1 female; 8 WT, 5 males and 3 females) were used for *in vivo* flat fiber photometry experiments. 3 mice (2 5xFAD, 2 males; 1 male WT) were used for *ex vivo* tapered fiber photometry experiments. 25 mice (14 5xFAD, 6 males and 8 females; 11 WT, 4 males and 7 females) were used for *in vivo* TF-based photometry experiments. Some mice were excluded from analysis due to recording or histological issues including an incomplete recording, inaccurate laser power estimations, or inaccurate registration of histological sections (**Supplementary Table 1**). No blinding or randomization was adopted, as well as analysis on sex differences due to this being a technical development study.

### *In vivo* flat fiber photometry

#### Photometry system

A flat fiber-based photometry system is described elsewhere (23). To adjust the system for Methoxy-X04 signals, the light from a 405 nm LED (M405L3, Thorlabs) was collimated by an aspheric lens (AL2520M-A, Thorlabs), passed a bandpass filter (FB405-10, Thorlabs), reflected off two dichroic mirrors (MD498 and DMLP425R, Thorlabs) and focused by an aspheric lens onto the multimode patch cable (400 μm core, 0.5 NA; MAF3L1, Thorlabs) and flat fiber (200 μm core, 0.50 NA; MAF3L1, Thorlabs). Emitted light passed back through the fiber and patch cable and was collimated by an aspheric lens, passed a dichroic mirror (DMLP425R, Thorlabs), and was redirected towards an emission filter by a broadband mirror (BB1-E02, Thorlabs). Filtered light was focused by an aspheric lens onto a photodetector (NewFocus 2151, Newport) for measurement. 440 nm and 550 nm emission filters (FB440-10 and FB550-10, Thorlabs) were used interchangeably by adding and removing a drop-in filter holder (DCP1, Thorlabs) between each measurement. A NIDAQ device (NI USB-6211, National Instruments) and custom LabVIEW software was used to control the LED and photodetector.

#### Photometry experiments

5xFAD+ and WT mice were used for *in vivo* flat fiber depth profile photometry experiments. 10 mg/kg of Methoxy-X04 (4920, Tocris) dissolved in 45% propylene glycol/45% 0.1 M phosphate buffered saline (PBS)/10% DMSO was administered intraperitoneally. On the next day, mice were anesthetized with urethane (1.5 g/kg) and placed in a stereotaxic frame (Model 963, KOPF Instruments). Body temperature was maintained at 37°C (ATC 100, World Precision Instruments) and eye gel (Hylonight or Viscotears) was applied throughout. Craniotomies were made over three sites (Site 1: AP +0.49 mm; ML 0.25 mm; Site 2: AP –1.79 mm, ML 1.50 mm; Site 3: AP –3.30 mm, ML 2.80 mm), which were covered with KWIK-SIL (World Precision Instruments). Throughout these procedures, the anesthetic level was maintained with isoflurane (1-1.5% at 0.8 L/min air flow).

Mice were moved to the recording set-up (SR-8N-S, Narishige) for depth profile measurements, where body temperature was maintained at 37°C (50-7212, Harvard Apparatus). For each craniotomy, KWIK-SIL was removed, and the flat fiber (200 μm core, 0.50 NA; MAF3L1, Thorlabs) was lowered to the brain surface by the substage and micromanipulator (SM-15M and SM-15, Narishige). A measurement was completed with both 440 and 550 nm emission filters, before repeating this process at increasing 100 μm depths until a maximum depth of 4000 μm was reached.

For each depth measurement, the LED went through 10 repetitions of 10 ms ON, 5 ms OFF, separated by a 10 ms baseline for light powers ranging from 0.1-1 mW/mm^2^. Data was collected at the photodetector and LED sync channel at 5000 Hz.

#### Histological analysis

Immediately after photometry experiments, mice were deeply anesthetized with lidocaine (2%) and pentobarbital (200 mg/ml) and transcardially perfused with PBS and 4% paraformaldehyde. Brain tissue was removed and post-fixated in the same fixative at 4°C overnight, and cryoprotected in 30% sucrose/0.1 M PBS for several days. Then, 100 μm coronal sections were prepared using a sliding microtome (SM2010R, Leica). As Methoxy-X04 was injected the day before brain retrieval, amyloid pathology was stained accordingly, and no other staining was necessary. Therefore, sections were washed in PBS (3 x 5 minutes), before rinsing in gelatin and mounting onto glass slides. Once dry, slides were sealed with cover slips and fluoromount solution (ThermoFisher Scientific Fluoromount G, Invitrogen). Brain sections anterior and posterior to and containing the flat fiber implant site were imaged. Images were acquired using an upright fluorescence microscope (Eclipse E600, Nikon) and CMOS camera (C11440-36U, Hamamatsu) at 4x magnification.

The post-processing of histological images is summarized in **Supplementary Figure 1**. Images were stitched automatically and the stitched images were registered to the Allen Mouse Brain Common Coordinate Framework using a modified version of a freely available software (AMaSiNe, https://github.com/vsnnlab/AMaSiNe) (28). Plaques were automatically detected as particles between 11-20 μm diameter. Coordinates of the flat fiber track were manually annotated across sections and estimated by using a linear regression model.

To estimate the plaque density, since the ipsilateral hemisphere was damaged by the flat fiber, the estimated fiber track was projected onto the contralateral hemisphere assuming that plaques are distributed symmetrically between hemispheres. To quantify plaques, a cylinder with a radius of 250 μm was generated along the projected fiber track and plaques within 250 μm from each depth measure were counted every 100 µm step. As virtually no plaque signals were detected in WT mice, randomly generated noise ranging between 0 and 1e-10 was added to histological quantification of both 5xFAD and WT mice without producing artificial trends. Plaque density was determined by calculating the plaque count per spherical volume at each quantified depth.

#### Photometry signal processing

Offline analysis was performed using custom MATLAB codes. For all analysis, 440 nm photometry data at 1 mW/mm^2^ was used. To obtain a depth profile, z-scored fluorescence at each depth was calculated by, 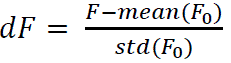, where *F* is the raw fluorescence and *F*_0_ is the mean fluorescence across 0-400 μm. A median filter was applied to smooth data (window size: 4). For comparison of photometry to histology signals, Spearman’s Rho correlation test was completed.

To classify photometry signals, each depth profile was Z-scored and the data matrix containing all samples (animals and recording sites) was constructed. Since signals were sampled at 41 depths for each penetration, principal component analysis (PCA) was applied to reduce the dimension. The first three PCs were used to train a support vector machine model to predict genotype. The training was done by taking the leave-one-out procedure and the genotype of the remaining sample was predicted by the trained model. The same procedures were repeated across all samples. To assess the performance of this classification, a confusion matrix was computed.

### *Ex vivo* tapered fiber photometry

#### Photometry system

A custom built two-photon laser scanning microscope described in detail in (29) was used to evaluate the capability of TFs to gather significant data from plaques-originated fluorescence. The microscope is equipped with three different imaging channels: an epi-fluorescence channel provided with a dichroic mirror (FF665-Di02, Semrock) and bandpass filter (FF01-520/70, Semrock), to acquire the reference image of the brain slices and the TF; a fiber-coupled channel provided with a bandpass filter (FF01-442/42-25, Semrock) and synchronized to the microscope scanner, to acquire the TF light collection field; and a wide-field channel provided with a bandpass filter (FBH450-40, Thorlabs) to acquire the TF illumination field on the sensor of a sCMOS camera (Orca Flash lite 4.0, Hamamatsu).

#### *Ex vivo* photometry experiments

To label amyloid pathology, Methoxy-X04 (10 mg/kg) (4920, Tocris) was injected intraperitoneally. The following day, brain samples were retrieved as described above. However, for this procedure, the sample did not undergo cryoprotection in sucrose and rather was stored in PBS. Then, 500 μm coronal sections were prepared using a vibratome (054018, Ted Pella Incorporated) and stored in PBS.

A brain slice containing the septal area was placed in the imaging plane of the custom-built two-photon laser scanning microscope and a TF (0.39 NA, 200 μm core, 225 μm cladding, 1.8 mm emission length, OptogeniX) was inserted into the slice so that the 1.8 mm optically active region of the taper was fully within the septal area.

Femtosecond pulsed laser light at 740 nm was used to elicit Methoxy-X04 fluorescence in brain slices through the microscope to measure the reference image and TF collection. Continuous wave 405 nm laser light (MDL-III-405-50-mW, CNI laser) was used to elicit Methoxy-X04 fluorescence in brain slices through the TF to measure the TF illumination field. Control on the angle at which light couples to the fiber was achieved by using a galvo mirror-based scanning system identical to the one employed for the *in vivo* experiments (described later in the text).

#### *Ex vivo* data analysis

Images were processed with FIJI software (30) and custom python scripts. Images acquired in the reference and fiber channels were intrinsically registered by syncing both PMTs with the laser scanning. Illumination fields acquired on a CCD camera were re-scaled and manually registered on the reference and fiber image. All images were preprocessed with background subtraction and thresholding before performing automating amyloid plaques localization and counting (Analyze Particles). The distribution of the plaques along the fiber axis was calculated on the images from the reference and fiber channels by computing the number of plaques centers falling within a circular ROI of radius 250 µm and centered on the fiber axis that was moved along the fiber at steps of 40 µm. The correlation between the quantified plaques in the reference image and fiber collection image was calculated using the Spearman’s rho correlation test.

The image stack of the illuminated areas of tissue against the input angles (galvo voltages) were pre-processed by background subtraction and normalized to the maximum value of the stack, to account for the dynamic in illumination density. For each, frame – corresponding to an input angle – a centroid of the illumination peak was calculated after subtracting an intensity baseline corresponding to the intensity of the terminal input angles, where a negligible amount of light is delivered in the tissue.

The photometry stack was then calculated by multiplying the illumination stack against the fiber channel image pixel by pixel, as described in (29). The simulated photometry intensities were calculated from the integral intensity in the photometry stack. Each value of photometry intensity was then attributed to a spatial location along the fiber corresponding to the centroid of the peak of the illumination profile for the corresponding input angle. This allowed to compare the photometry intensity profiles against the plaque density profile in the reference image (intensity scaled by dividing by the maximum value). The correlation between the photometry intensity profiles and the intensity scaled plaque density in the reference image was calculated using the Spearman’s rho correlation test.

### *In vivo* tapered fiber photometry

#### Photometry system

A 405 nm laser (MDL-III-405-50-mW, CI90055, CNI lasers or Cobolt 06-01 Series, Hubner Photonics) delivered excitation light. A portion of this light beam was reflected towards a photodetector (PDA25K-EC, Thorlabs) by a glass slide to monitor laser stability. The rest passed broadband mirrors before being focused onto a galvo mirror (GVS001, Thorlabs) by an aspheric lens (AC254-050-A-ML, Thorlabs). The galvo mirror controlled the angle of reflected light and therefore, the angle light coupled into the tapered fiber (0.39 NA, 200 μm core, 225 μm cladding, 1.8 mm emission length, OptogeniX). Reflected light was collimated by an aspheric lens (AC254-050-A-ML, Thorlabs), passed through a dichroic mirror (DMSP425T, Thorlabs) before being focused by an aspheric lens (AC254-030-A-ML, Thorlabs) onto a multimode patch cable (200 μm core, 0.39° NA) and an implanted tapered fiber. Emitted light passed back through the tapered fiber and patch cable and was collimated by an aspheric lens and reflected off a dichroic mirror (DMSP425T, Thorlabs) towards a dichroic mirror (DMSP490R, Thorlabs) which separated 440 nm and 550 nm light beams, with use of emission filters (FB440-10 and FB550-10, Thorlabs). Each beam was then focused onto the photodetector by an aspheric lens (AC254-030-A-ML, Thorlabs) for measurement. A NIDAQ device (NI USB-6343, National Instruments) and custom LabVIEW software was used to control the laser, galvo mirror and photodetectors.

#### Calibration and photometry protocols

For equalizing the light power across the tapered fiber, the fiber was placed in uniform Methoxy-X04 solution (0.1 mM) and underwent an illumination protocol. The laser was activated in cycles of 10 ms ON, 5 ms OFF at 60 and 80 μW for 5 repetitions. In parallel with laser ON periods, the galvo mirror was driven from –1 to 4.5 V, remaining at 0 V during the OFF period. Data was collected at 5000 Hz. The fluorescence at each galvo measure was used to calculate the required power for uniform fluorescence across the tapered fiber. For each galvo measure, the mean fluorescence was transformed to a relative required power by, 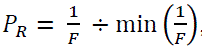, where *P*_*R*_ is the required power and *F* is the fluorescence.

#### Photometry experiments

5xFAD+ and WT mice were used. Mice were anaesthetized with isoflurane (5% for induction and 1-1.5% for maintenance with 0.8 L/min airflow). Mice were placed in a stereotaxic frame (Model 963, KOPF Instruments), reflexes were monitored, body temperature was maintained at 37°C (ATC 100, World Precision Instruments) and eye gel (Hylonight or Viscotears) was applied throughout. Naropin (8 mg/kg) was administered to the surgical area subcutaneously, while Vetergesic (0.1 mg/kg), Rimadyl (20 mg/kg) and saline (0.3 ml, FKE1323, Baxter) were administered subcutaneously to the rump. Skull screws (418-7123, RS Components) were implanted in the skull: two in the front (AP +1.50 mm, ML ±1.00 mm), one over the right hemisphere (AP –1.00 mm, ML +4 mm), one over the left hemisphere (AP –3.00 mm, ML –4.00 mm) and two in the back (AP –2.00 mm from lambda, ML ±2.00 mm) as anchors. The tapered fiber was implanted at the target site (AP –3.50 or –3.60 mm, ML 2.25 or 2.80 mm, DV 2.50 or 2.00 mm) before being secured with KWIK-KAST (World Precision Instruments) and layers of superglue (918-6872, RS-Pro) and dental cement (Kemdent). Dental cement was applied across the skull and skin edges. After 5 days of recovery, mice were habituated to handling and being tethered to the photometry system. While scruffed, they were attached to the patch cable via a mating sleeve (ADAL1-5, Thorlabs) and placed into a recording chamber (46.5 x 21.5 x 20.5 cm). Recordings included a baseline measurement with no Methoxy-X04 for 5 hours. 24-hour later a 5-hour measurement was completed, with Methoxy-X04 (10 mg/kg) (4920, Tocris) being injected intraperitoneally at 30 minutes. 24 hours later, the fluorescence was monitored for a further two hours. The laser was activated in cycles of 10 ms ON, 5 ms OFF across 4 powers ranging from 60-140 μW. In parallel with laser ON periods, the galvo mirror was driven from –1 to 4.5 V, remaining at 0 V during the OFF period. This protocol was repeated 3-5 times at a sampling interval of 60 or 300 s. Data was collected at 5000 Hz. After several recordings, mice were culled for histological analysis.

#### Histological analysis

Brain sections (50 μm) were prepared as described above. As Methoxy-X04 was injected several days before brain retrieval, amyloid pathology will be stained accordingly, but sections were co-stained with another plaque marker as a back-up. To identify the TF track, sections were co-stained with a microglial marker. After washes in PBS (3 x 5 mins), sections were incubated in 10% normal goat serum in 0.3% Triton X in PBS (PBST) for 1 hour. Then, sections underwent primary antibody incubation with anti-Iba1 antibody (1:1000 in 3% normal goat serum in PBST; Abcam, cat. no. ab178846) overnight at 4°C. After washes in PBS (3 x 5 mins), a 2-hour secondary antibody incubation with Alexa Flour 594 (1:1000 in 3% normal goat serum in PBST; ThermoFisher, cat. no. A-11005) was completed. Sections were washed in PBS (3 x 5 mins) before co-staining with Thioflavin-S (0.01% in PBS; Sigma-Aldrich, cat. no. T1892) for 15 minutes. Sections were washed in PBS (3 x 5 minutes), before rinsing in gelatin and mounting onto glass slides. Once dry, slides were sealed with cover slips and fluoromount solution (ThermoFisher Scientific Fluoromount G, Invitrogen). Brain sections were imaged as described above.

Image processing was the same as described above. Plaque quantification was the same as described above but was completed at 41 steps every 39.5 μm (galvo measure resolution). Also, to account for overestimations occurring during the estimation of the TF penetration track, we shifted the penetration track up 600 μm **(Supplementary Figure 2A)**.

#### Photometry signal processing

Offline analysis was performed using custom MATLAB codes. To estimate Methoxy-X04-related fluorescent signals, the dynamics of the autofluorescence was predicted based on the signals without Methoxy-X04 administration (Day 0). To this end, a linear model was established at each galvo scanning level as *AF* = *a*(*t*) + *b*, where *AF* was predicted autofluorescence, *t* was the time samples and *a* and *b* are the model coefficients, which were estimated by two-fold cross-validation. Methoxy-X04 signals were estimated at each Galvo scanning level as *F*_*m*_ = *F* − *AF*, where *F*_*m*_ was the estimated Methoxy-X04 signals and *F* was raw fluorescent signals. Measurements from 0-4.5 V galvo scanning levels were taken in 5-minute time bins across recordings. The depths containing the minimum and maximum median signal from 30-240 minutes were z-scored by, 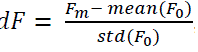, where *dF* was the z-scored fluorescence, *F*_*m*_ was estimated Methoxy-X04 signals and *F*_0_was the mean fluorescence in the first 25 minutes of recording. Data was smoothed using a moving median filter. For all analysis, 440 nm photometry data at 120 μW was used.

### Statistical analysis

All statistical analysis was performed using MATLAB otherwise stated. A Kolmogorov–Smirnov test was completed to determine the normality of the data. Accordingly, in Figures 1, 2 and 3, correlation analysis was completed using the Spearman’s Rho test. A two-sample t-test was performed in Figures 1 and 3 to compare correlation coefficients. In Figure 3, the maximum and minimum fluorescent profiles for 5xFAD and WT were tested with a Wilcoxon signed-rank test. All data was shown as mean ± standard error of mean unless otherwise stated.

**Figure 1.**
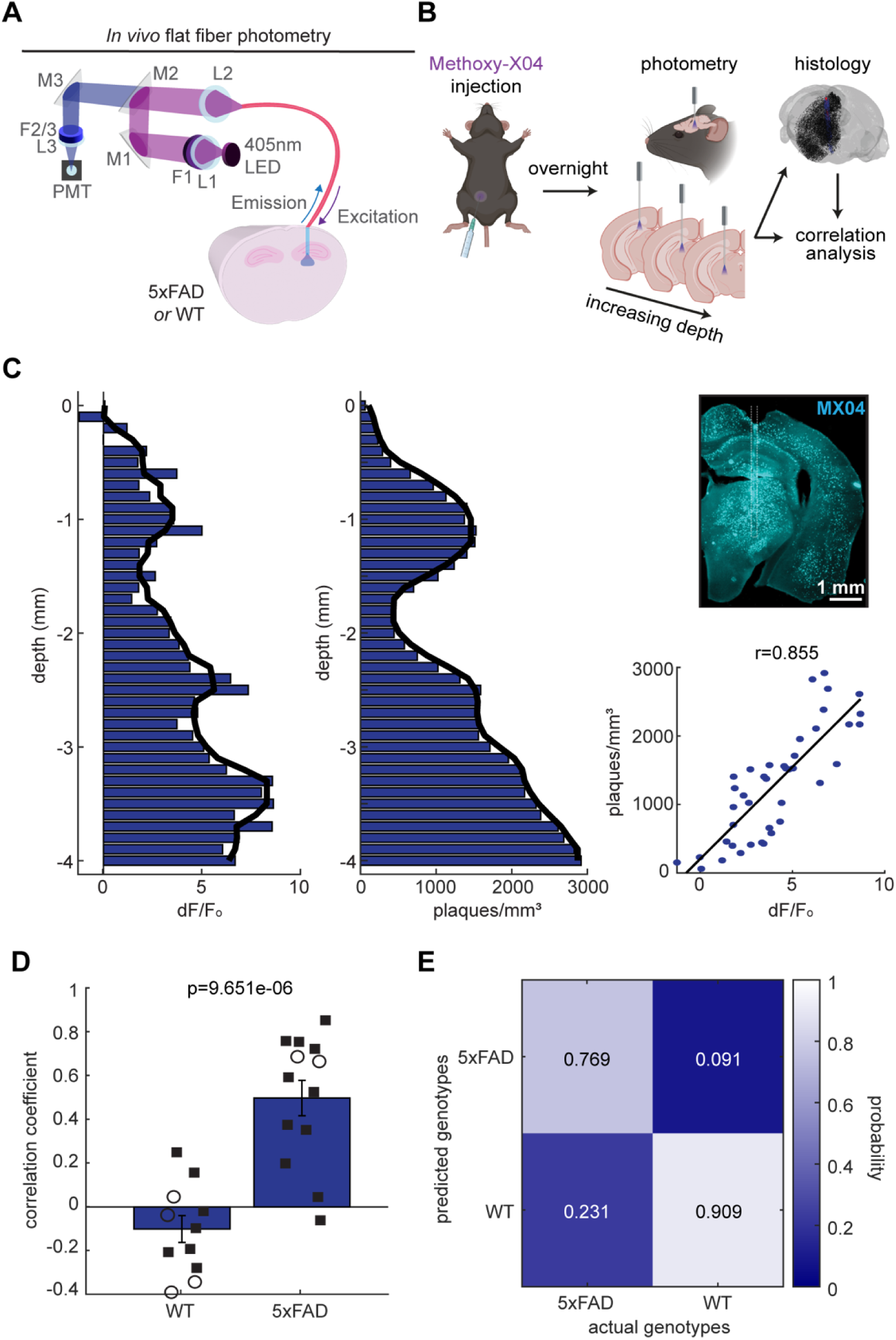
Flat fiber-based photometry realizes fluorescent signals reflective of amyloid pathology in 5xFAD mice. (**A**) Flat fiber-based photometry system allows 405 nm LED light to pass the flat fiber implanted in either 5xFAD or WT mouse brain. Emitted fluorescence light passes back through the flat fiber before being filtered to cut out excitation light and being focused onto a photodetector (PMT). The light path was controlled by a series of mirrors (M), filters (F) and lenses (L). (**B**) Schematic of the experimental design. Mice were injected with Methoxy-X04 the day before being terminally anesthetized for photometry recordings. After, brain tissue was retrieved for histology where photometry and histology signals underwent correlation analysis. (**C**) *Left,* Example *in vivo* fluorescent and post-mortem histology depth profiles. Solid line shows the median smoothed signal (window size: 4). *Top right*, Coronal brain slice showing the flat fiber penetration track surrounded by Methoxy-X04-stained amyloid pathology. White dashed line shows the penetration track. Scale: 1 mm. *Bottom right*, Correlation analysis comparing photometry and histology depth profiles (r, Spearman’s Rho). Black line shows a fitted linear regression. (**D**) Summary correlation coefficients across three different implant sites (two-sample t-test). 5xFAD, n = 13 recordings from 6 mice. WT, n = 11 recordings from 7 mice. Squares, males; Circles, females. (**E**) Classification performance based on *in vivo* photometry-based depth profiles.

**Figure 2.**
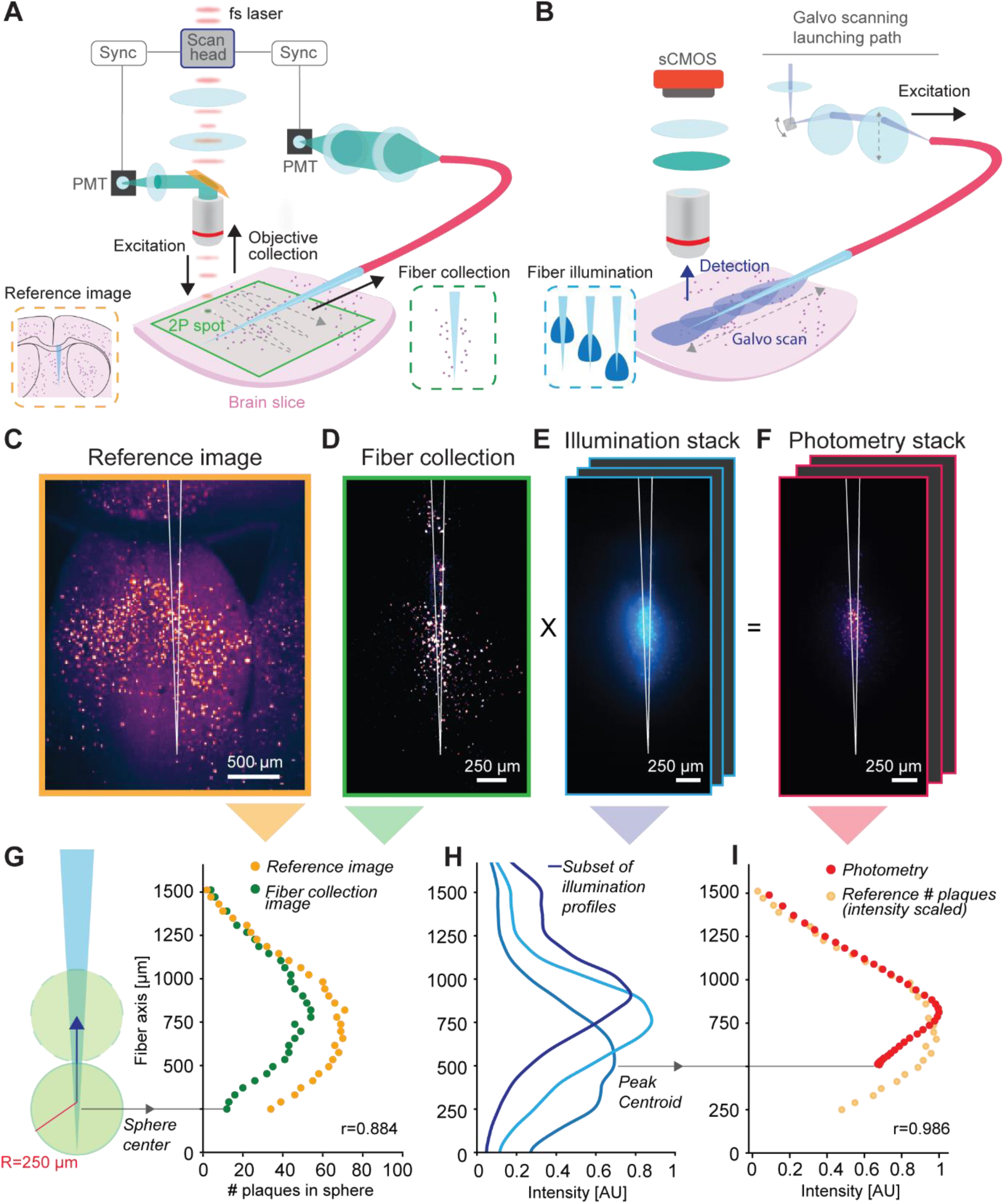
*Ex vivo* TF-based photometry of amyloid plaques. (**A**) Schematic of the two-photon imaging system used to acquire co-registered reference images and fiber collection images. (**B**) Schematic of the angle-selective light injection system and imaging system to acquire images of the illumination profiles obtained by the TF in brain tissue. (**C**) Reference image of the amyloid plaques around the implanted TF outlined in white. (**D**) Fiber collection image of the amyloid plaques, with the fiber profile outlined in white. (**E**) Example of illumination image collected varying the injection angle in the TF. (**F**) Calculated photometry stack, obtained as a pixel-by-pixel multiplication of the illumination stack versus the collection image. (**G**) Number of plaques in a moving region of interest modelled as a circle with 250 µm radius for the reference (orange dots) and collection (green dots) image (r, Spearman’s Rho). (**H**) Example of a subset of the illumination profiles measured along the TF for each frame in the illumination stack corresponding to an input galvo voltage (**E**). The centroid of the peak illumination intensity for each input voltage is used as a reference position of the photometry intensity. (**I**) Simulated photometry intensity profile extracted from the photometry stack; the photometry intensity profile (red dots) is consistent with the distribution of plaques extracted from the reference image (orange dots, rescaled for comparison) (Spearman’s Rho correlation test).

**Figure 3.**
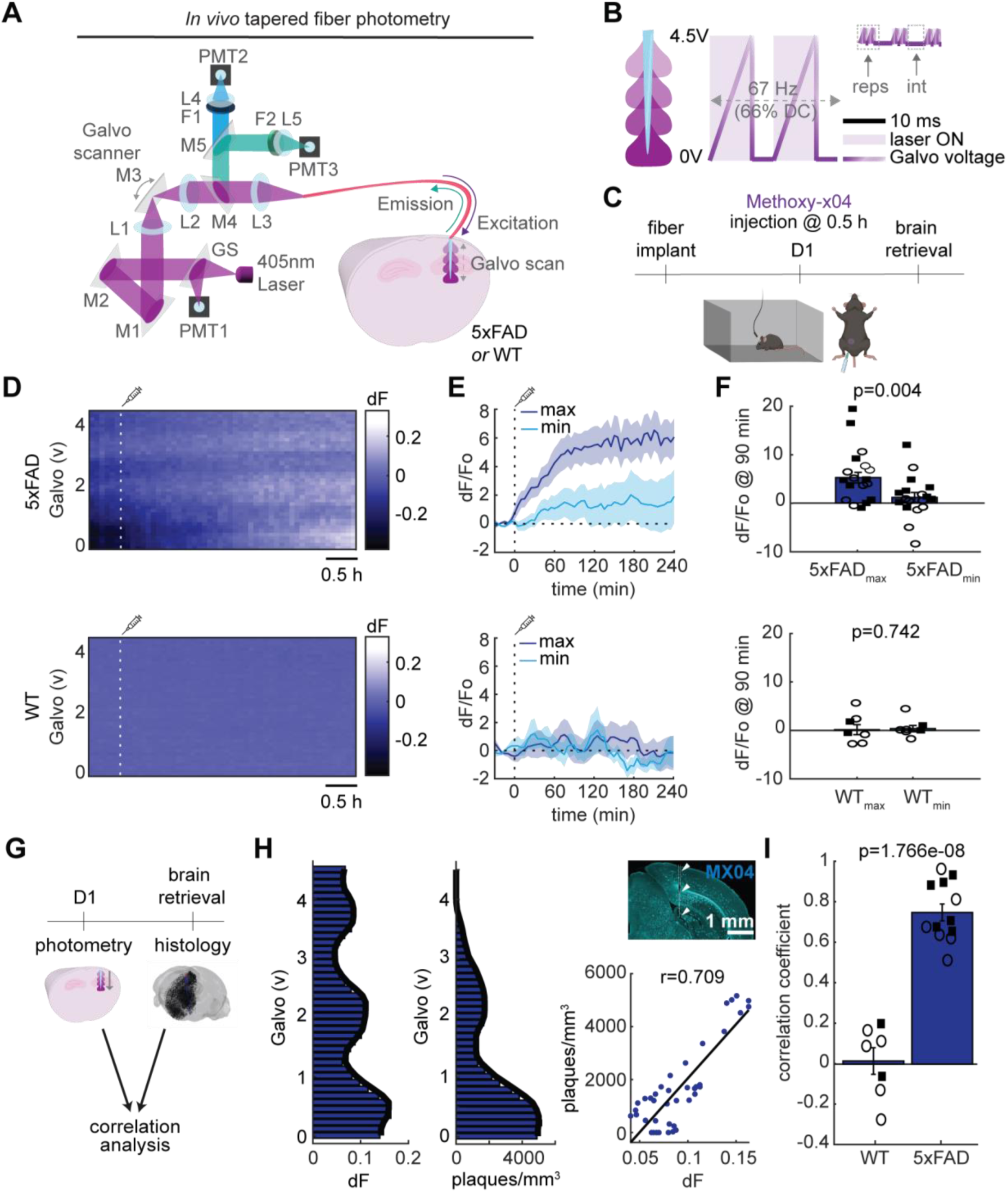
TF-based photometry of amyloid plaques in freely behaving 5xFAD mice. (**A**) TF-based photometry system allows 405 nm laser light to pass the tapered fiber implanted in either 5xFAD or WT mouse brain. Emitted light passes back through the tapered fiber before being focused onto a PMT for each emission wavelength. Light propagation along the tapered fiber was controlled by a galvo mirror, allowing increasing depths to be achieved in one complete galvo scan. The light path was controlled by a series of glass slides (GS), mirrors (M), filters (F) and lenses (L). (**B**) Illumination protocol. The 67 Hz cycle was repeated (reps) 3 or 5 times, at a 1-or 5-minutes interval (int). (**C**) Schematic of the experimental design. Methoxy-X04 was injected (i.p.) after 0.5 hours on day 1 (D1) of the recording session. (**D**) Example heatmaps of *in vivo* fluorescence depth profiles, shown over galvo voltage, for the D1 recording session. White line shows Methoxy-X04 injection. (**E**) Depths (galvo voltages) providing the maximum and minimum fluorescence changes for 5xFAD and WT mice. 5xFAD, n = 18 recordings from 12 mice; WT, n = 8 recordings from 7 mice. Some mice were recorded from more than once. (**F**) Depth-specific fluorescence change at 90-minutes (Wilcoxon signed-rank test). Squares, males; Circles, females. (**G**) Comparisons between photometry and histology signals. (**H**) *Left,* Example *in vivo* fluorescent and post-mortem histology depth profiles. Solid line shows the median smoothed signal (window size: 4). *Top right*, Methoxy-X04-stained coronal section showing the tapered fiber penetration track (white dashed line and arrow heads). *Bottom right*, Correlation between photometry and histology depth profiles (r, Spearman’s Rho). (**I**) Summary correlation coefficients (two-sample t-test). 5xFAD, n = 12 mice; WT, n = 7 mice. Squares, males; Circles, females.

## Results

### Flat fiber-based photometry to monitor amyloid pathology in anesthetized 5xFAD mice

As a proof-of-concept experiment, we firstly examined if conventional flat fiber-based photometry (**Fig. 1A**) allows monitoring of plaque signals across multiple depths *in vivo*. To this end, Methoxy-X04 was injected a day before experiments in either 5xFAD mice (n = 6) or their littermate wild-type (WT) controls (n = 7). Under urethane anesthesia, we performed fiber photometry at increasing depths by gradually lowering a flat fiber into the brain (**Fig. 1B**). As we lowered the flat fiber, we observed variable fluorescent signals along the penetration track in a 5xFAD mouse (**Fig. 1C**). After photometry experiments, we performed histological analysis to reconstruct the depth profile of plaque density along the penetration track (**Fig. 1C**). We took the depth profile from the contralateral hemisphere to estimate signals in the intact brain. We confirmed a significant positive correlation between *in vivo* fluorescent and post-mortem histological depth profiles (r = 0.855, p < 0.0001, Spearman’s Rho) (**Fig. 1C**). In contrast, signals in a control animal showed lower variability. Since the control animal did not form plaques, no correlation was found (**Supplementary Figure 3A**). We repeated the same experiments across three different sites across animals to find statistically significant positive correlations in 5xFAD mice (0.499 ± 0.081) compared to control mice (–0.101 ± 0.061) (t(22) = 5.70, p < 0.0001, two-sample t-test) (**Fig. 1D**). These results indicate that *in vivo* photometry signals reflect plaque signals at depth.

We also examined whether photometry-based depth profiles alone can discriminate genotypes (i.e., 5xFAD or WT). To this end, we took a machine learning approach. After reducing the dimension to 3 from 41 data points across the penetration by performing principal component analysis, we trained a support vector machine model by leaving one sample out. Then we tested the model to assess if the remaining sample was accurately classified (**Fig. 1E**). The overall performance was 83.3%: out of 13 5xFAD recordings, ten were properly classified (76.9% hit) whereas out of 11, ten WT recordings were correctly rejected (90.9%). These results indicate that *in vivo* photometry signals alone are capable of classifying genotypes.

### *Ex vivo* evaluation of tapered fiber-based photometry

Given the success of the flat fiber-based approach, we sought to develop a method to monitor amyloid pathology across brain regions in freely behaving animals. To this end, we adopted tapered optical fiber (TF) technology (20, 31). Before using TFs *in vivo*, we examined if a TF allowed depth-resolved photometry of amyloid plaques *ex vivo* in a controlled relevant environment (**Fig. 2**). To this end, we injected Methoxy-X04 into a 5xFAD mouse and extracted the brain tissue on the following day. We then inserted a TF (0.39NA, 200 µm core, 225 µm cladding, 1.8 mm emission length) in a 500 µm thick coronal brain slice containing the septal area and used a two-photon scanning microscope equipped with an epi-fluorescence channel and a fiber-coupled channel (**Fig. 2A**) (29, 32) to simultaneously collect a reference image of the brain slice (**Fig. 2C**) and a measurement of the TF collection field (**Fig. 2D**). To determine if the fiber collection field was large enough to gather a significant amount of plaques-originated fluorescence, we compared the number of plaques enclosed inside a 250 µm radius sphere with the center moving along the TF axis on these two images (**Fig. 2G**): despite the fiber collecting a lower number of plaques than the epi-fluorescence channel, mainly due to the decay in collection efficiency while moving radially farther from the fiber axis, the two curves are consistent (r = 0.884, p < 0.0001, Spearman’s Rho).

With the TF in the same position, we delivered 405 nm light across different positions along the TF to the brain slice while changing the laser input angle with a galvanometer mirror based scanning system (**Fig. 2B**) (20); for each input angle we collected an image by using a wide-field path on the two-photon microscope to measure the light illumination fields produced by the TF at every input voltage of the galvo (**Fig. 2E**). This allowed us to estimate the portion of the fiber collection field involved in fluorescence excitation for different input angles by multiplication of the fiber collection field and the fiber illumination stack (**Fig. 2F**), obtaining a stack of images representing the photometry fields. The images in the illumination stack were also used to locate the centroid of the area of tissue recruited at every input angle using the position of the centroid of the intensity profile taken along the fiber (**Fig. 2H**). An *ex vivo* simulated photometry signal was then calculated as the sum of all the pixel intensities in the photometry stack, and each datapoint was positioned according to the corresponding illumination profile. We compared the curve obtained in this way with a re-scaled version of the plaques count in the reference image and found a significant correlation between the two measurements (r = 0.986, p < 0.0001, Spearman’s Rho) (**Fig. 2I**). These results suggest that TF-based photometry can be used to monitor amyloid pathology across brain regions *in vivo*.

### Depth-resolved photometry to monitor amyloid pathology in freely behaving 5xFAD mice

To monitor plaque signals in a freely behaving condition, we constructed a photometry system with a 405 nm laser and galvo mirror (**Fig. 3A**). We configured the galvo mirror to scan signals along the TF (∼1.6 mm) 3 or 5 times at 66 Hz (67% duty cycle). We repeated this scanning every 1 or 5 minutes to monitor fluorescence signals along the TF (**Fig. 3B**). Since amyloid plaques densely appear in the subiculum in this mouse model (27, 33), we chronically implanted a TF into the subiculum and surrounding regions (5xFAD, n = 18 recordings from 12 mice; WT, n = 8 recordings from 7 mice).

Our typical recording schedule consisted of a baseline recording without Methoxy-X04 (Day 0) and a recording with a Methoxy-X04 injection (Day 1) (**Fig. 3C**). On Day 1, after Methoxy-X04 injection, the signals gradually increased in 5xFAD mice with variable signal intensity across depth, but not in WT mice (**Fig. 3D**).

To assess if the increased fluorescent signals reflect depth-resolved plaque distribution, we conducted two analyses: first, we compared the fluorescent dynamics on Day 1 at the galvo levels which provided the maximum and minimum signal intensities (**Fig. 3E**). We clearly observed differential intensity changes approximately 1.5 hours after Methoxy-X04 injections in 5xFAD mice (**Fig. 3E**). On the other hand, we did not see such differential changes in WT. Specifically, at 1.5 hours post-injection we see a depth-resolved change in intensity in 5xFAD mice (p = 0.004, Wilcoxon signed-rank test) but not in WT mice (p = 0.742, Wilcoxon signed-rank test) (**Fig. 3F**).

Second, to correlate the depth profiles between *in vivo* photometry and post-mortem histology, we compared the photometry depth profile from 210-240 minutes to the reconstructed histology plaque density along the penetration track (**Fig. 3G**). Similar to the acute experiments (**Fig. 2**), we took the depth profile from the contralateral intact hemisphere to estimate histological signals. We confirmed a significant positive correlation between *in vivo* fluorescent and post-mortem histological depth profiles (r = 0.709, p < 0.0001, Spearman’s Rho) (**Fig. 3H**). In contrast, since the WT animal did not form plaques, no correlation was found (**Supplementary Figure 3B**). Overall, we found significant positive correlations in 5xFAD mice (0.750 ± 0.042) compared to WT mice (0.015 ± 0.066) (t(17) = 9.911, p < 0.0001, two-sample t-test). We also assessed how robust the correlation between *in vivo* and histological signals was against a range of parameters (i.e., the TF tip estimation and light dispersion radius, see also the Materials and Methods section). We found that the high correlation in 5xFAD mice was generally preserved across parameter values (**Supplementary Figure 2**). Finally, we observed less tissue damage with a TF compared to a flat fiber (**Supplementary Figure 4**), consistent with a previous report (34). Thus, we established a TF-based photometry approach to monitor amyloid plaque signals across brain regions, in a freely behaving condition.

## Discussion

In the present study, we reported a novel fiber photometry-based approach to monitor amyloid plaque signals *in vivo*. First, using a conventional flat fiber-based approach, we confirmed that amyloid plaque signals can be monitored across depth, under anesthesia, *in vivo* (**Fig. 1**). Second, we assessed if TFs allow depth-resolved photometry for plaque signals *ex vivo* (**Fig. 2**). After confirming the feasibility *ex vivo*, we validated our TF-based approach in a freely behaving condition (**Fig. 3**).

### Comparisons to other methods

Amyloid pathology in mouse models has been assessed by various methods, yet postmortem histological analysis remains the gold standard. While positron emission tomography (PET) scan or microdialysis has been used to detect amyloid plaques (10, 35), the former is not readily available for many researchers and the latter cannot provide spatially resolved signals. Optical approaches have been sought before: although two-photon microscopy has been used (13–15), it is challenging to obtain signals in deep brain structures. While optoacoustic tomography allows monitoring of brain-wide plaque signals (16), the imaging needs to be done under anesthesia. Our approach offers a unique, novel solution to monitor depth-resolved plaque signals in a freely behaving condition.

While fiber photometry has been adopted for a wide range of experiments in neuroscience (17, 18), our approach has expanded its potential to assess molecular pathology in an AD mouse model *in vivo*. We demonstrated that fiber photometry can be used without a genetic strategy, rather with injection of a blood-brain barrier permeable fluorescent dye. This enhances the adaptability of this technology for various cellular and molecular processes, without the requirement of a genetically encoded construct. Also, compared to flat fiber-based photometry (**Fig. 1**), our TF-based photometry offers unique advantages. In addition to less invasiveness, we demonstrated that TF-based technology enables us to obtain depth-resolved plaque signals in a freely behaving condition for the first time. While TF technology has been applied in various ways (20, 26, 36, 37), the present study opens a new avenue for preclinical applications of TF-based technology in neurodegenerative disease mouse models.

### Limitations of the study

Despite the success of this novel approach, we are aware of several limitations. First, although Methoxy-X04 has been widely used, the excitation spectrum is blue-shifted (i.e., ∼405 nm). This requires a dedicated laser to excite Methoxy-X04. Since a ∼470 nm light source is commonly used in the field, the adoption of the reported methodology requires extra investment in an optical setup. On the other hand, combining another wavelength of light will be straightforward. For example, it may be interesting to apply an optogenetic approach by expressing red-shifted opsins to modulate neural activity (38).

Second, although a TF allows depth-resolved photometry, the length of the taper can be up to 3 mm (31). With the current technology, it is not practical to monitor plaque distribution across the brain. Like in the present study, an approach targeting several regions is suitable. Multi-fiber arrays (24, 39) may be an alternative approach to monitor plaque signals across brain regions in a freely behaving condition. At the same time, recent advances in deep brain spectroscopy using tapered fibers also open an interesting perspective to complement our method with reporter free detection (40).

Third, like all other photometry-based approaches, our approach cannot resolve individual plaque signals. Despite this limitation, we confirmed that the depth profile of photometry signals is correlated with histological signals across experimental conditions. It may be interesting to explore advanced optical approaches, such as an endo-microscope (41), to monitor individual plaques although the issue with the absorption spectrum of Methoxy-X04 needs to be addressed.

## Conclusion

Our novel fiber photometry approach allows us to monitor amyloid plaque signals at depth in a freely behaving condition. Our results demonstrate the novel potential of *in vivo* fiber photometry, which has been widely used in the neuroscience community. Our TF-based photometry approach also opens new avenues for monitoring plaque signals to optimize therapeutic approaches and to develop novel intervention strategies for AD in a preclinical setting.

## Acknowledgements

This work was supported by Medical Research Council (MR/V033964/1 and MR/Y004051/1 to S.S.), Alzheimer’s Research UK (PPG2023B-019 to S.S.), the European Union’s Horizon 2020 (H2020-ICT, DEEPER, 101016787 to K.M., M.D.V., Fe.P. and S.S.; MINIG, 101125498 to M.D.V. and Fe.P.), and the PARD 2024 from the University of Padua to Fi.P. K.M. was supported through an RAEng Chair in Emerging Technologies. C.M., M.D.V. and Fe.P. acknowledge funding from the Project ‘RAISE (Robotics and AI for Socio-economic Empowerment), (code ECS00000035) funded by European Union – NextGenerationEU PNRR MUR – M4C2 – Investimento 1.5 – Avviso ‘Ecosistemi dell’Innovazione’ CUP J33C22001220001.

## Author contributions

N.B. and S.S. conceived *in vivo* experiments. N.B. and N.M. developed the optical systems for *in vivo* experiments. N.B. performed all *in vivo* experiments and data analysis. N.B. and J.F. performed histological experiments. N.B., Fi.P., M.P., and S.S. conceived *ex vivo* experiments. Fi.P., M.P. and C.M. performed *ex vivo* experiments and data analysis. K.M., M.D.V., Fe.P. and S.S. supervised the work. N.B., Fi.P., M.P. and S.S. wrote the original manuscript. N.B., Fi.P., J.F., C.M., K.M., M.D.V., Fe.P. and S.S. reviewed and edited the manuscript.

## Supplementary Tables

**Supplementary Table 1.** Information about the mice used throughout this study. The experiment each mouse was used for is listed as flat fiber (FF), *ex vivo* (ExV) or tapered fiber (TF). Mice that were excluded and the appropriate reason is noted.

| ID | Genotype | Sex | Age @ experiment start (mo) | Excluded? | Reason | Experiment type | Figure no. |
| --- | --- | --- | --- | --- | --- | --- | --- |
| 1 | + | F | 6.9 |  |  | FF | 1 |
| 2 | + | M | 8.5 |  |  | FF | 1 |
| 3 | + | M | 8.4 |  |  | FF | 1 |
| 4 | + | M | 6.8 | Y | incomplete recording | FF |  |
| 5 | + | M | 7.6 |  |  | FF | 1 |
| 6 | + | M | 6.9 |  |  | FF | 1 |
| 7 | + | M | 7.1 |  |  | FF | 1 |
| 8 | - | F | 8.0 |  |  | FF | 1 |
| 9 | - | F | 6.9 |  |  | FF | 1 |
| 10 | - | F | 8.4 |  |  | FF | 1 |
| 11 | - | M | 8.4 |  |  | FF | 1 |
| 12 | - | M | 6.9 | Y | incomplete recording | FF |  |
| 13 | - | M | 7.0 |  |  | FF | 1 |
| 14 | - | M | 7.1 |  |  | FF | 1 |
| 15 | - | M | 7.1 |  |  | FF | 1 |
| 16 | + | M | 9.1 |  |  | ExV | 2 |
| 17 | + | M | 9.1 |  |  | ExV | 2 |
| 18 | - | M | 9.1 |  |  | ExV | 2 |
| 19 | + | F | 4.3 | Y | incorrect laser power estimation | TF |  |
| 20 | + | F | 4.8 |  |  | TF | 3 |
| 21 | + | F | 6.0 |  |  | TF | 3 |
| 22 | + | F | 3.0 |  |  | TF | 3 |
| 23 | + | F | 3.3 |  |  | TF | 3 |
| 24 | + | F | 8.0 | Y | histology registration error | TF |  |
| 25 | + | F | 3.5 |  |  | TF | 3 |
| 26 | + | F | 6.8 |  |  | TF | 3 |
| 27 | - | F | 4.6 | Y | incorrect laser power estimation | TF |  |
| 28 | - | F | 4.9 |  |  | TF | 3 |
| 29 | - | F | 6.0 |  |  | TF | 3 |
| 30 | - | F | 2.9 |  |  | TF | 3 |
| 31 | - | F | 3.3 |  |  | TF | 3 |
| 32 | - | F | 8.0 |  |  | TF | 3 |
| 33 | - | F | 3.5 |  |  | TF | 3 |
| 34 | + | M | 4.9 |  |  | TF | 3 |
| 35 | + | M | 6.0 |  |  | TF | 3 |
| 36 | + | M | 3.0 |  |  | TF | 3 |
| 37 | + | M | 3.9 |  |  | TF | 3 |
| 38 | + | M | 3.6 |  |  | TF | 3 |
| 39 | + | M | 6.8 |  |  | TF | 3 |
| 40 | - | M | 4.7 | Y | incorrect laser power estimation | TF |  |
| 41 | - | M | 3.0 |  |  | TF | 3 |
| 42 | - | M | 3.9 | Y | histology registration error | TF |  |
| 43 | - | M | 5.7 |  |  | TF | 3 |

## Supplementary Figures

**Supplementary figure 1.**
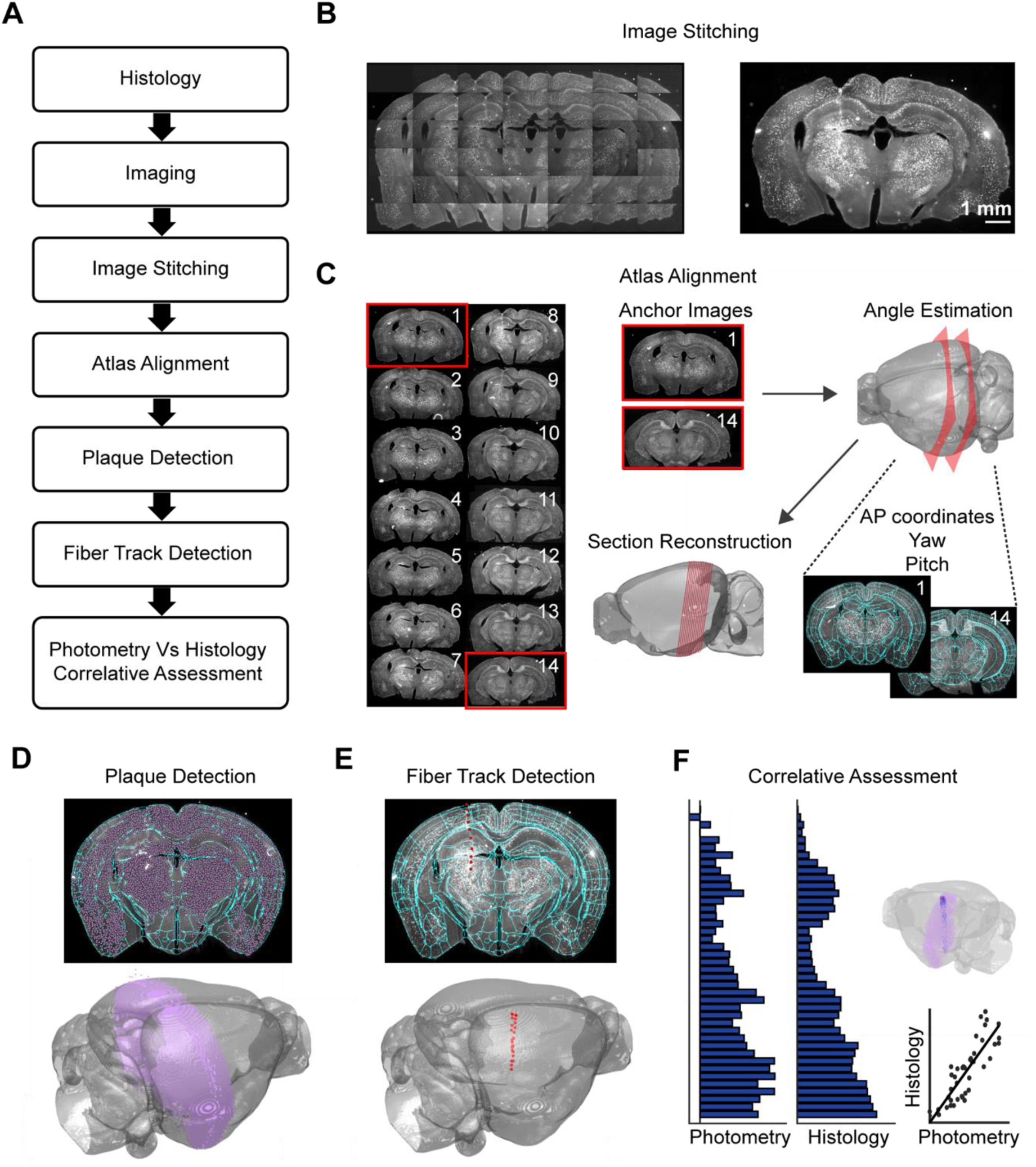
Protocol for reconstruction, registration and quantification of Methoxy-X04 stained histological sections. (**A**) Flow chart of the protocol. (**B**) Image stitching of all original images (*left*) to the final stitched image (*right*). Scale bar: 1 mm. (**C**) The first and final stitched image (*shown with red border*) are anchor images used for atlas alignment. Anchor images are used to determine several alignment parameters before all image slices are reconstructed onto the atlas. (**D**) *Top*, Automated plaque detection on an aligned brain slice. *Bottom,* Detected plaques on all aligned brain slices, visualized on a whole brain. Purple signals represent a detected plaque. (**E**) *Top*, Manual fiber track detection on an aligned brain slice. *Bottom,* Detected fiber track on all aligned brain slices, visualized on a whole brain. Red signals represent the manually annotated fiber track. (**F**) Plaques within 250-μm of the manually annotated fiber track (*red*), on the contralateral hemisphere are quantified. Blue signals represent quantified plaques. Correlation analysis was completed on photometry and histology depth profiles.

**Supplementary figure 2.**
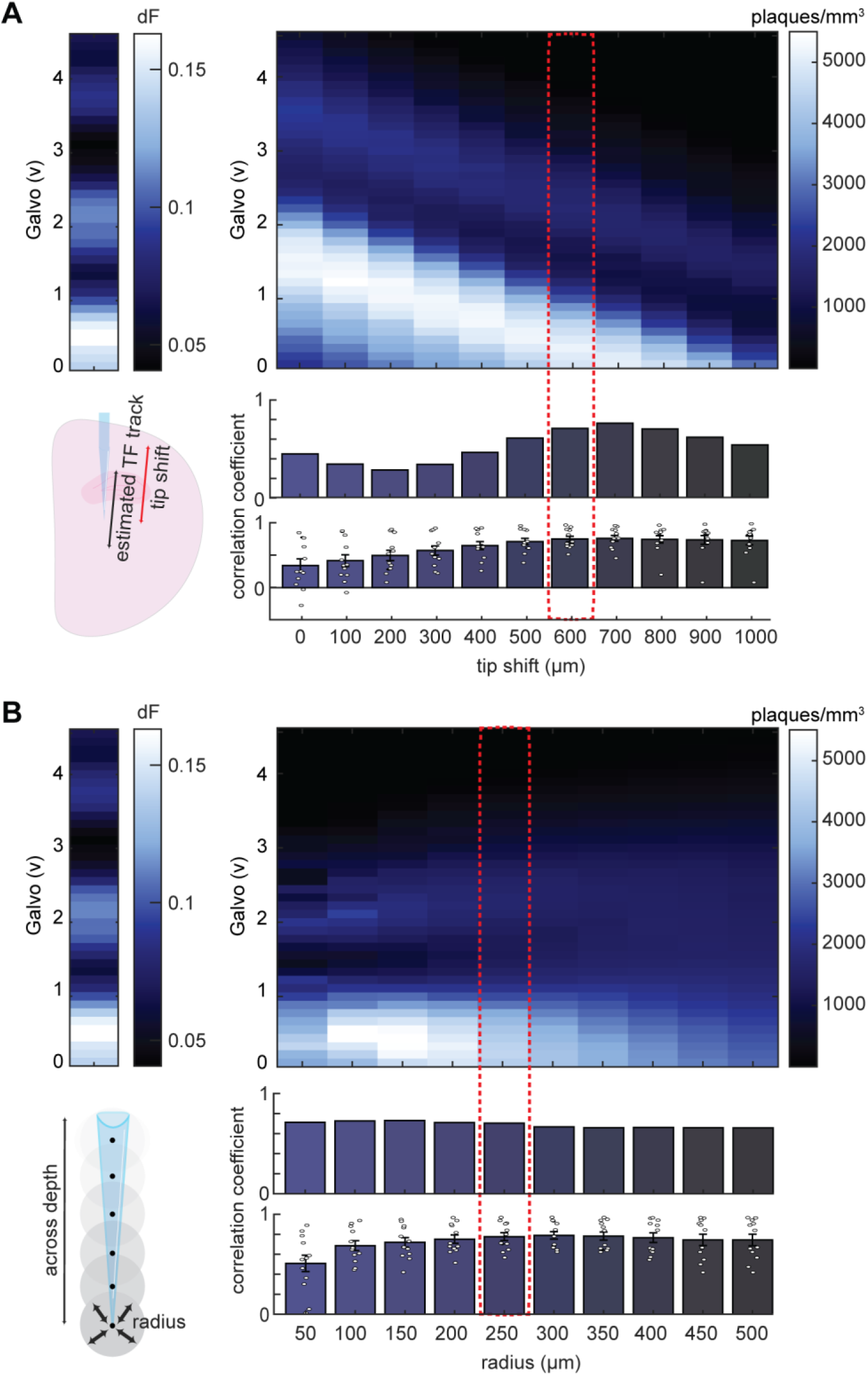
Systemic assessment of the histological quantification parameters. (**A**) *Bottom left,* TFs are minimally invasive which makes it challenging to accurately identify the TF tip on histological sections. Often, this led to an error in the estimated penetration track for TF implants. This error could be corrected by shifting the TF penetration track up several hundred micrometers. *Top left,* Heatmap of the mean fluorescent signals from 210-240 minutes on day 1 of the TF recording. *Top right*, Heatmap of the histology depth profiles with TF tip shifts from 0 to 1000 μm. *Bottom right*, Top panel shows the correlation coefficient when comparing the photometry and histology depth profiles for this example. Bottom panel shows the correlation coefficient across all mice. Red dashed line shows a tip shift of 600 μm provides the best correction. (**B**) *Bottom left,* The radius of the sphere represents the distance threshold for quantified plaques used at each galvo measure. *Top left,* Heatmap of the mean fluorescent signals from 210-240 minutes on day 1 of the TF recording. *Top right*, Heatmap of the histology depth profiles with radius from 50 to 500 μm. *Bottom right*, Top panel shows the correlation coefficient when comparing the photometry and histology depth profiles for this example. Bottom panel shows the correlation coefficient across all mice. Red dashed line shows a radius of 250 μm provides the best reflection of light propagation from the TF.

**Supplementary figure 3.**
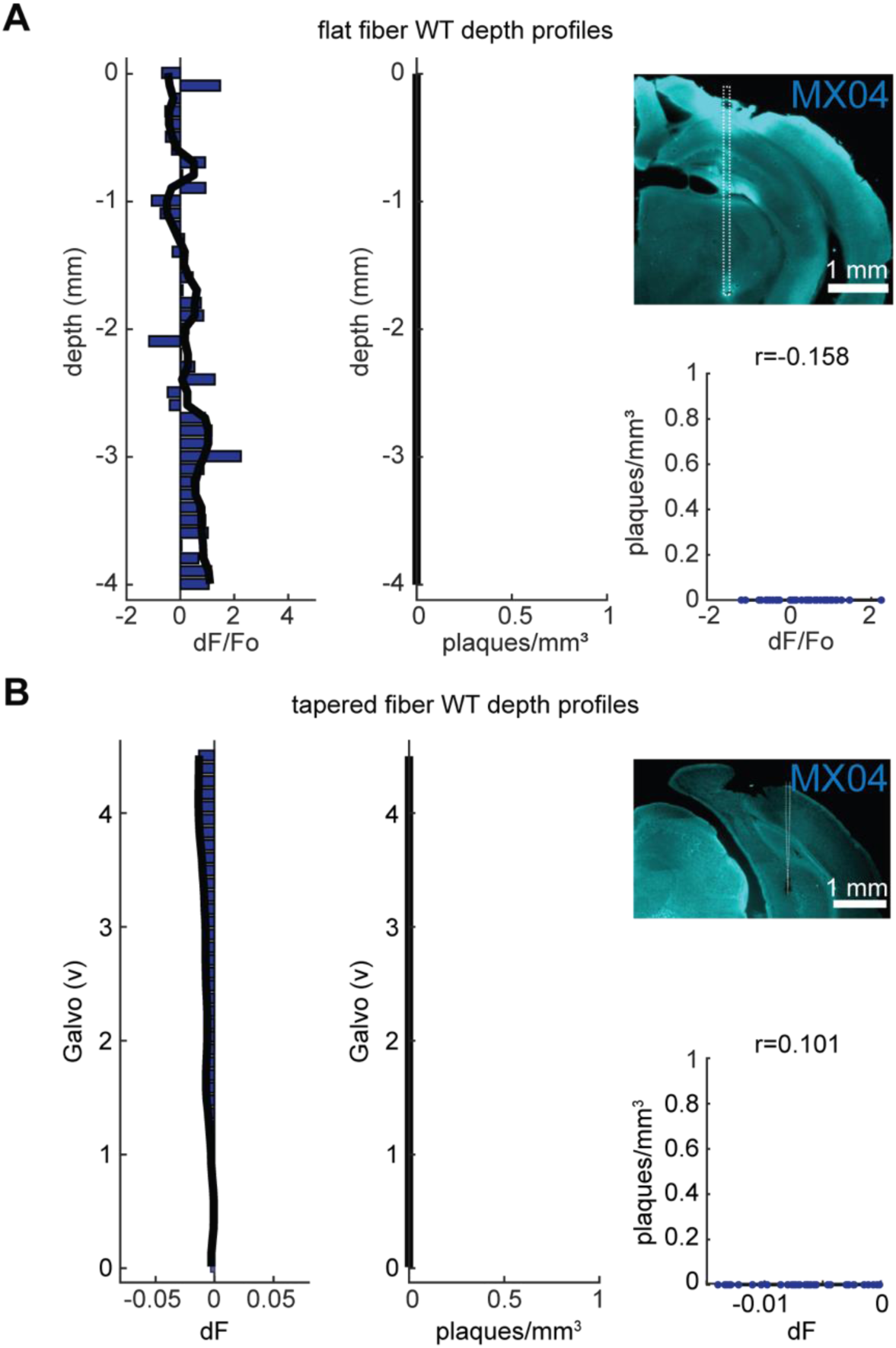
**Fiber photometry and histological assessment of Methoxy-X04 signals in WT mice**. Examples of flat fiber-based (**A**) and TF-based (**B**) experiments. *Left,* Example *in vivo* fluorescent and post-mortem histology depth profiles. Solid line shows the median smoothed signal (window size: 4). *Top right*, Coronal brain slice showing the fiber penetration track (white dashed line). Scale: 1 mm. *Bottom right*, Correlation analysis comparing photometry and histology depth profiles (Spearman’s Rho correlation test). Black line shows a fitted linear regression.

**Supplementary figure 4.**
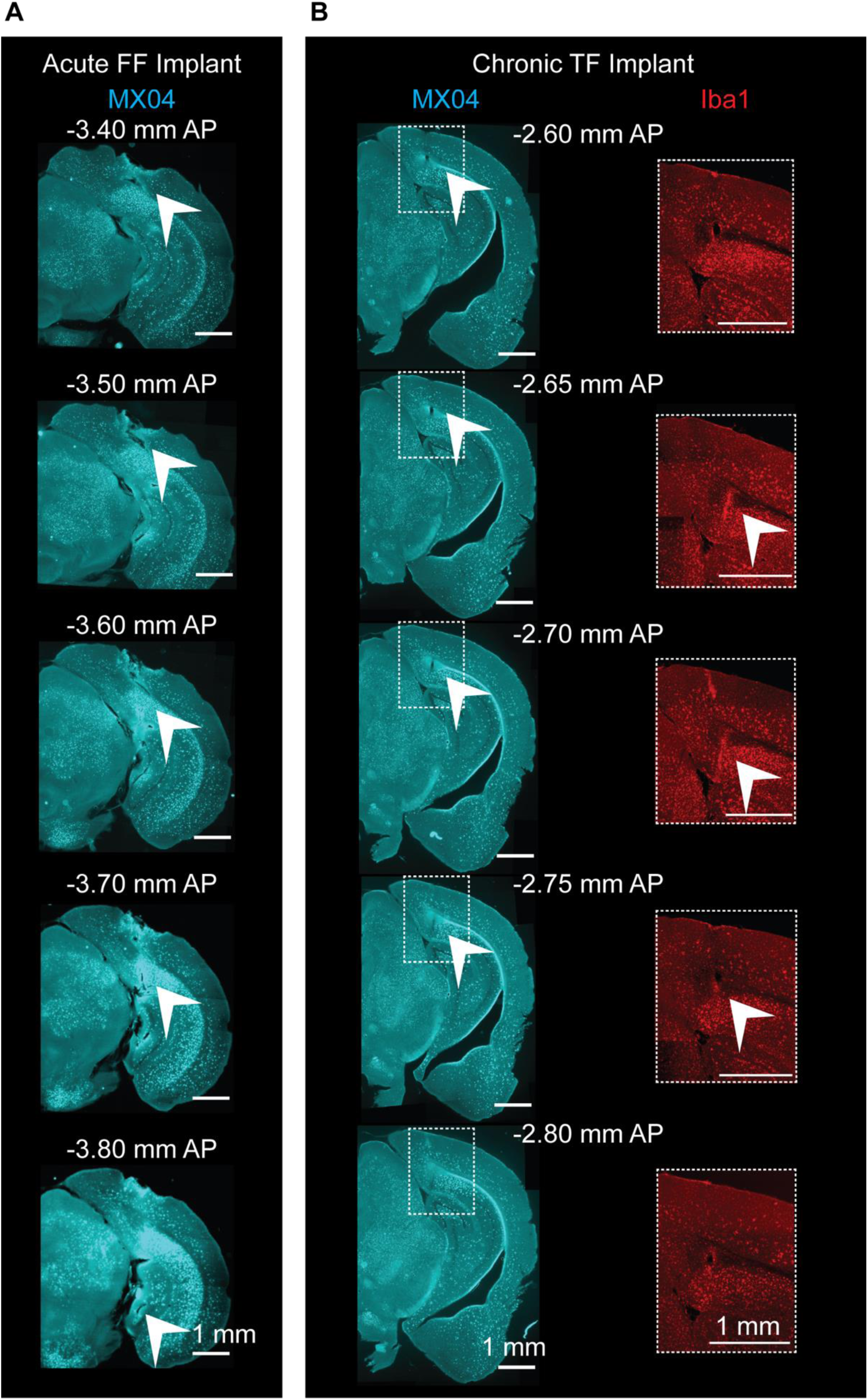
**Assessment of damage and microgliosis after acute FF and chronic TF implantation**. (**A**) Coronal Methoxy-x04 stained sections across the AP axis show tissue damage over a large area after an acute FF implant. White arrows show tissue damage. (**B**) Coronal Methoxy-x04 stained sections across the AP axis show less tissue damage and gliosis (zoomed inset in white dashed line) over a small area. White arrows show tissue damage and areas of microgliosis. Scale: 1 mm.

